# Inflammatory disease microbiomes share a functional pathogenicity predicted by C-reactive protein

**DOI:** 10.1101/2025.01.14.633015

**Authors:** Graham J. Britton, Lorenza Bartu, Ilaria Mogno, Alice Chen-Liaw, Tamar Plitt, Drew Helmus, Gerold Bongers, India Brough, Paula Colmenero, Lilian H. Lam, Samuel J. Bullers, Frank Penkava, Pamela Reyes-Mercedes, Jonathan Braun, Jonathan P. Jacobs, A. Nicole Desch, Dirk Gevers, Sheri Simmons, Andrew Filer, Peter C. Taylor, Paul Bowness, Curtis Huttenhower, Dan Littman, Marla C. Dubinsky, Karim Raza, Stephanie K. Tankou, Jeremiah J. Faith

## Abstract

We examine disease-specific and cross-disease functions of the human gut microbiome by colonizing germ-free mice, at risk for inflammatory arthritis, colitis, or neuroinflammation, with over 100 human fecal microbiomes from subjects with rheumatoid arthritis, ankylosing spondylitis, multiple sclerosis, ulcerative colitis, Crohn’s disease, or colorectal cancer. We find common inflammatory phenotypes driven by microbiomes from individuals with intestinal inflammation or inflammatory arthritis, as well as distinct functions specific to microbiomes from multiple sclerosis patients. Inflammatory disease in mice colonized with human microbiomes correlated with systemic inflammation, measured by C-reactive protein, in the human donors. These cross-disease patterns of human microbiome pathogenicity mirror features of the inflammatory diseases, including therapeutic targets and the presence or absence of systemic inflammation, suggesting shared and disease-specific mechanisms by which the microbiome is shaped and drives pathogenic inflammatory responses.

## INTRODUCTION

Inflammation is a hallmark of disease – crucial for infection defense and tissue repair but pathogenic if unresolved. Immune-mediated inflammatory diseases (IMIDs), including inflammatory bowel disease (IBD), rheumatoid arthritis (RA) and multiple sclerosis (MS), are characterized by maladapted inflammation in specific tissues. Inflammation can also manifest in the absence of a specific condition; sub-clinical or ‘low-grade’ inflammation (LGI) is a major independent risk factor for cardiovascular disease, type-2-diabetes and cancers (*1-3*). Furthermore, inflammation in the intestine in the context of ulcerative colitis is a major risk-factor for colorectal cancer (*4*). The composition of the intestinal microbiome is frequently altered in patients with IMIDs (*5-9*) and inflammation-associated cancers (*10, 11*), and disruption of the host-microbiome interface has been implicated in LGI (*12-15*). These results motivate the hypothesis that the intestinal microbiome contributes to the initiation or exacerbation of inflammation in these diverse and burdensome conditions (*5, 16, 17*).

Studies of microbiome composition in human disease typically focus on individual diseases, while cross-disease comparisons have been limited in scale, or use metanalysis of multiple datasets where technical variation can mask biological effects (*18-22*). However, a survey of studies that focus on individual diseases reveals that changes that are observed in microbiome structure and composition in human IMIDs tend to be similar, even amongst diseases that affect tissues distal from the gut. These changes, commonly referred to as dysbiosis, are characterized by a reduction in the diversity and density of the intestinal microbiome, as well as characteristic shifts in the abundance of specific taxa (*5-10, 23-25*). However, the directionality of the interaction between pathogenic inflammation and the intestinal microbiome is unclear, as host factors including diet, therapeutics and inflammation itself may alter the microbiome with no direct impact on pathology in the patient (*26*).

Evidence for a direct causative role of microbiome is well defined in ulcerative colitis (UC), where fecal microbiota transplant (FMT) has shown some clinical efficacy (*27-32*). Beyond this, the best evidence for a direct role for the microbiome in the etiology of IMIDs has come from animal models. By colonizing formerly germ-free mice with the gut microbiota of humans with or without IMIDs, several studies of various scale have demonstrated that microbiome transfer is sufficient to recapitulate features of the human condition in mouse models (*33, 34*). For example, transfer of microbiomes from humans with RA, MS or IBD to germ free mice can exacerbate disease in mouse models of arthritis (*35*), experimental autoimmune encephalomyelitis (EAE) (*36, 37*) and colitis (*38*), respectively. However, given shared characteristics of the microbiome across IMIDs, it is unknown if the effects in animal models are disease-specific or if they collectively represent a pan-disease proinflammatory shift in the microbiome, irrespective of the human IMID or animal model. This question is of fundamental importance for understanding the relationship between the microbiome and inflammation in etiology, pathogenesis and as a therapeutic target.

In this study, we colonize germ free mice with over 100 human microbiomes from donors with a spectrum of inflammatory diseases or inflammation-associated cancer and assess the influence of these microbiomes on three mouse models of inflammatory disease. We find that different human microbiomes dramatically modulate disease severity in mice susceptible to colitis, arthritis or neuroinflammation (EAE). Microbiomes collected from humans with MS exacerbate EAE but not colitis or arthritis. In contrast, microbiomes from humans with IBD or inflammatory arthritis exacerbate disease in mice with colitis or arthritis, but not EAE. Thus, we demonstrate that whereas microbiomes from humans with MS are functionally distinct from other tested IMID, microbiomes from humans with IBD, RA or ankylosing spondylitis (AS) are functionally similar. These observations parallel the underlying pathobiology and therapeutic targets in these conditions. Finally, we observe a relationship between systemic inflammation in the human microbiome donor and inflammation in mice colonized with that human’s intestinal microbiome. Together, these data reveal new insights into the contribution of the microbiome to inflammatory disease.

## RESULTS

We assembled a collection of over 100 human microbiomes including 70 samples from individuals diagnosed with a spectrum of IMIDs: Crohn’s disease (CD), ulcerative colitis (UC), RA, AS, MS, and inflammation-associated cancer (treatment-naïve colorectal cancer; CRC). IMIDs stools were matched with microbiomes from 30 healthy individuals collected as part of the same cohort studies from which we obtained the IMID samples (Table 1).

### In contrast to other IMID microbiomes, microbiomes from humans with multiple sclerosis exacerbate EAE

Experimental autoimmune encephalomyelitis (EAE) is a mouse model of central nervous system inflammation that is commonly used to model aspects of multiple sclerosis. Disease is dependent on an intact microbiome, and microbiomes from humans with multiple sclerosis have been shown to exacerbate disease symptoms in mice, with the largest study including 5 MS microbiomes (*36, 37*). We colonized groups of germ-free C57Bl/6 mice with one of 57 different fecal microbiomes (a total of 306 mice, median = 5 mice/microbiota) and induced EAE in these animals three weeks later. We find that different microbiomes have a significant influence on EAE severity (p<1 × 10^-6^, ANOVA; Figure 1A). On average, mice colonized with microbiomes from humans with MS experienced more severe EAE symptoms than mice colonized with healthy donor microbiomes, including higher peak disease scores and greater total disease burden (p=0.01 and p=0.03, respectively; ANOVA with Dunnett’s post-test; Figure 1B and C). Microbiomes from humans with AS, RA, UC, CD or CRC did not exacerbate EAE relative to healthy donor microbiomes (p>0.4), and disease in mice with these IMID microbiomes was less severe than mice with microbiomes from humans with MS (Peak disease; p=0.06) (Figure 1C). Among the human fecal donors with MS, neither donor disease severity (EDSS score) nor disease duration correlated with EAE severity in mice colonized with donor microbiomes (Fig S1).

**Fig. 1:**
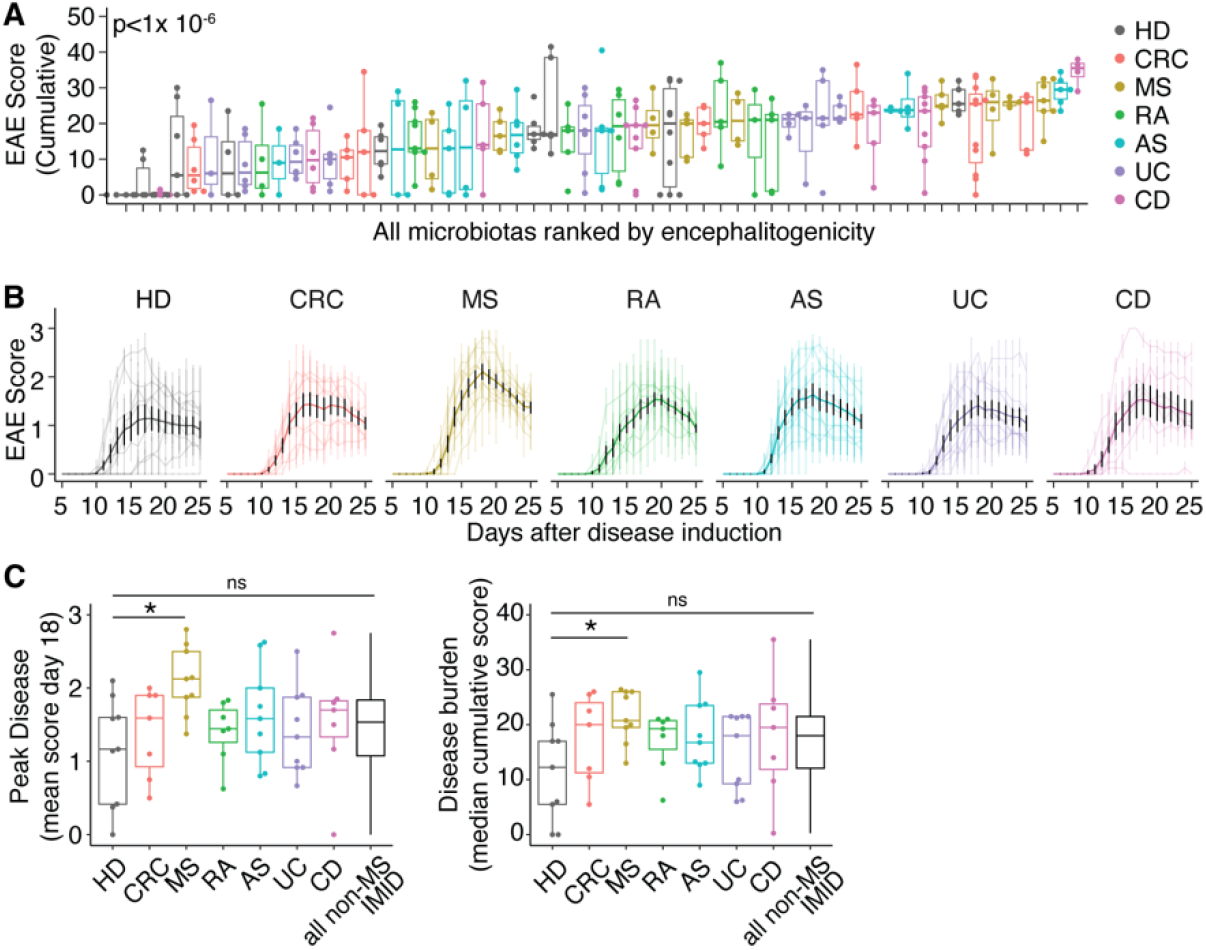
In contrast to other IMID microbiomes, microbiomes from humans with multiple sclerosis exacerbate EAE. **(A)**, EAE severity in mice colonized with one of 56 different human microbiomes (median cumulative EAE scores for each mouse). P value calculated by ANOVA. **(B)**, EAE scores over time of mice colonized with different human microbiomes. Lighter colored lines show median data of mice colonized with each microbiome; darker lines show the weighted mean EAE scores +/- SEM of mice colonized with microbiomes from humans with the indicated disease. **(C)**, Peak EAE severity (median score at day 18 after induction) and EAE disease burden (cumulative disease score day 0-25) of groups of mice colonized with different human microbiomes. *p<0.05, **p<0.01, Dunnett’s test with BH correction comparing each disease group to HD group. Non-MS IMID refers to data from all donors with AS, RA, CD or UC. In all panels, N human microbiomes of each disease tested in n mice: AS, N=8, n=41; CD, N=7, n=43; CRC, N=7, n=42; HD, N=9, n=55; MS, N=9, n=43; RA, N=7, n=37; UC, N=9, n=45. Boxplots show median +/- IQR in the style of Tukey.

### Microbiomes from humans with diverse IMID enhance arthritis and colitis severity in susceptible mice

The SKG mouse carries a mutation in the TCR-proximal signaling protein Zap70 that enforces selection of an exaggeratedly self-reactive TCR repertoire (*39*). In specific pathogen free (SPF) SKG mice on the BALB/c background, administration of a fungal adjuvant is sufficient to trigger multi-tissue autoinflammatory disease, most profoundly affecting the peripheral and spinal joints and the small intestine, mimicking aspects of ankylosing spondylitis, rheumatoid arthritis and Crohn’s disease (*40*). We colonized germ free SKG mice with a subset of the IMID, CRC and healthy donor microbiomes (36 microbiomes into a total of 186 mice, median 5 mice/microbiota) and induced arthritis-like disease through injection of fungal β-glucan curdlan. We observed significant inter-microbiome variation in the severity of disease in mice colonized with different human microbiomes, as measured by joint swelling, loss in body mass and symptom scoring (p<1 × 10^-6^, ANOVA; Figure 2A). A previous study reported that a pool of three microbiomes from donors with RA exacerbated disease in SKG mice compared to a pool of healthy donor microbiomes (*35*). Using individual, non-pooled microbiomes, we find that microbiomes from humans with RA or AS experience greater joint swelling, increased weight loss and more severe symptom scores relative to mice harboring healthy donor microbiomes (Figure 2B and C).

**Fig. 2:**
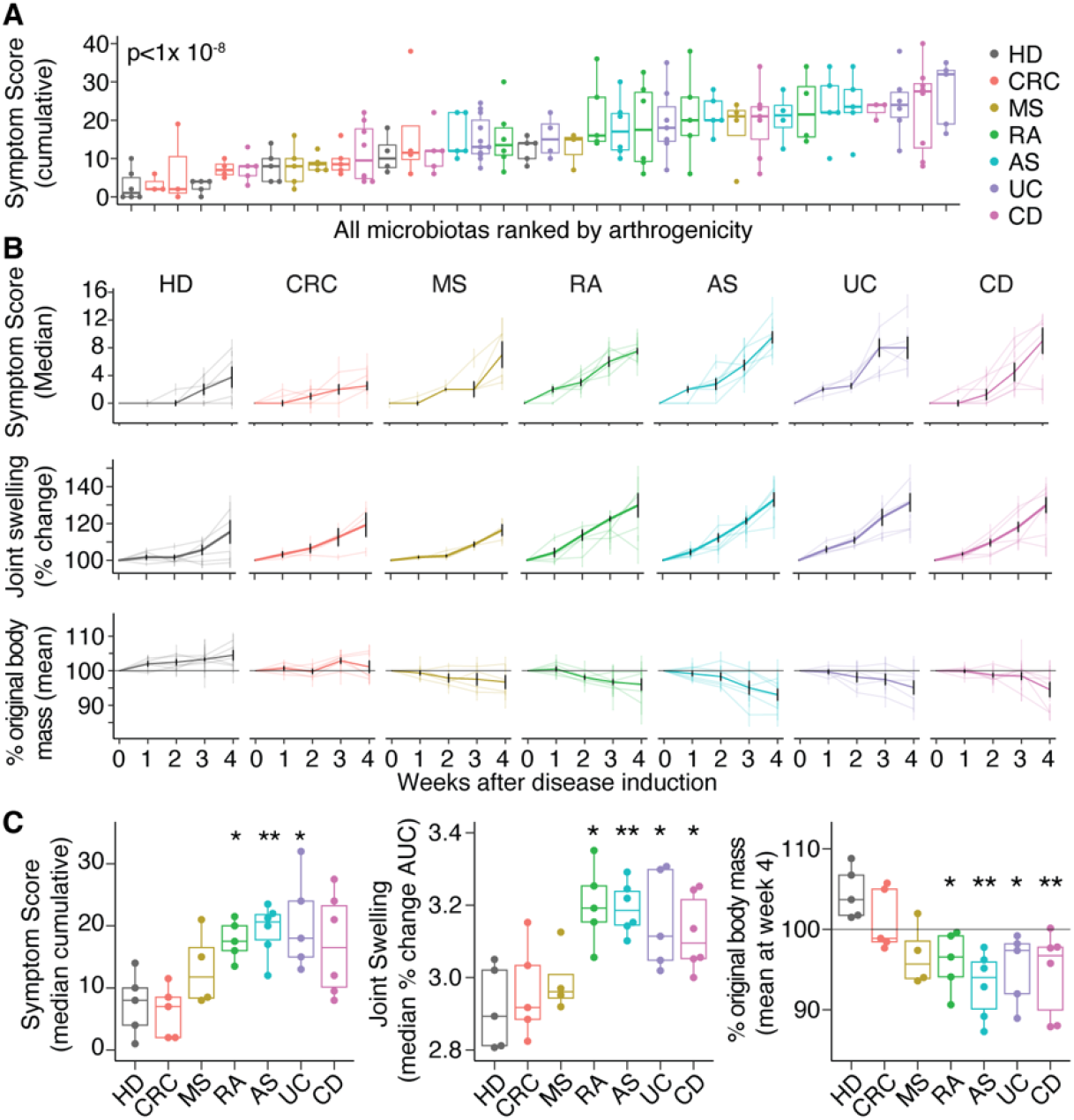
Microbiomes from humans with diverse IMID enhance arthritis severity in susceptible mice. **(A)**, Arthritis severity in SKG mice colonized with one of 35 different human microbiomes (points shows the cumulative symptom score of individual mice). Colors indicate the health status of the microbiome donor. P value calculated by ANOVA. **(B)**, Arthritis symptom scores, joint swelling (percentage change in joint dimensions from baseline) and change in body mass over time following disease induction in groups of mice colonized with different human microbiomes. Lighter colored lines show median (symptom scores) or mean (joint swelling and body mass) data of groups of mice colonized with each microbiome; darker lines show the median +/- SEM (symptom scores) or mean +/- SEM (joint swelling and body mass) of mice colonized with microbiomes from humans with the indicated disease. **(C)**, Median symptom scores (cumulative), joint swelling (area under the curve; AUC) and body mass change of mice colonized with microbiomes from each disease type. *p<0.05, **p<0.01, Dunnett’s test comparing disease group to HD group. In all panels, N human microbiomes of each disease tested in n mice: AS, N=5, n=30; CD, N=5, n=36; CRC, N=5, n=19; HD, N=5, n=25; MS, N=5, n=17; RA, N=5, n=26; UC, N=5, n=33. Boxplots show median +/- IQR in the style of Tukey.

Interestingly, microbiomes from humans with either UC or CD also exacerbated disease in SKG mice relative to healthy donor microbiomes. Microbiomes from humans with CRC or MS did not exacerbate joint swelling, symptom scores or weight loss when compared to healthy donor microbiomes (p>0.1 for all comparisons; Figure 2B and C). As previously reported (*41*), we observe more severe disease in female SKG mice as compared to male mice. IBD and inflammatory arthritis microbiomes exacerbated arthritis in SKG mice irrespective of sex (Figure S2).

We have previously demonstrated that microbiomes from humans with IBD exacerbate disease in a mouse model of colitis (*38, 42*). To assess if this is a property specific to IBD microbiomes, we introduced each of the IMID, CRC and healthy control microbiomes into groups of germ free C57Bl/6 Rag1^-/-^ mice (99 microbiomes in a total of 391 mice, median 3 mice/microbiota) and induced colitis-like disease in these animals by the adoptive transfer of naïve CD4^+^ T cells obtained from sex-matched SPF mice. The 99 microbiomes we tested induced a wide spectrum of disease severity, as assessed by the primary measure of weight loss (Figure 3A). Importantly, we observed no significant difference in weight loss between groups of healthy donor microbiotas coming from different cohorts (Figure S3A). Consistent with previous studies, microbiomes from humans with CD or UC exacerbated disease in these mice relative to microbiomes from healthy individuals as measured by loss in body mass, histological scoring of colon tissues and induction of the fecal inflammatory biomarker lipocalin-2 (LCN2) (*42, 43*) (Figure 3B and C, Figure S3B-D). Interestingly, microbiomes from humans with AS or RA also exacerbated loss in body mass relative to healthy donor microbiotas (Figure 3B and C, Figure S3B-D). The average disease severity induced by AS and RA microbiomes was not significantly different from CD or UC (p>0.1 for all comparisons). Microbiomes from humans with CRC or MS did not significantly exacerbate colitis relative to microbiomes from healthy donors (p>0.8). Consistent with our previous studies using the T cell transfer model of colitis (*38*), loss in body mass was strongly correlated with direct measures of intestinal inflammation (scoring of colon histopathology and fecal LCN2 concentrations) (Figure S3E).

**Fig. 3:**
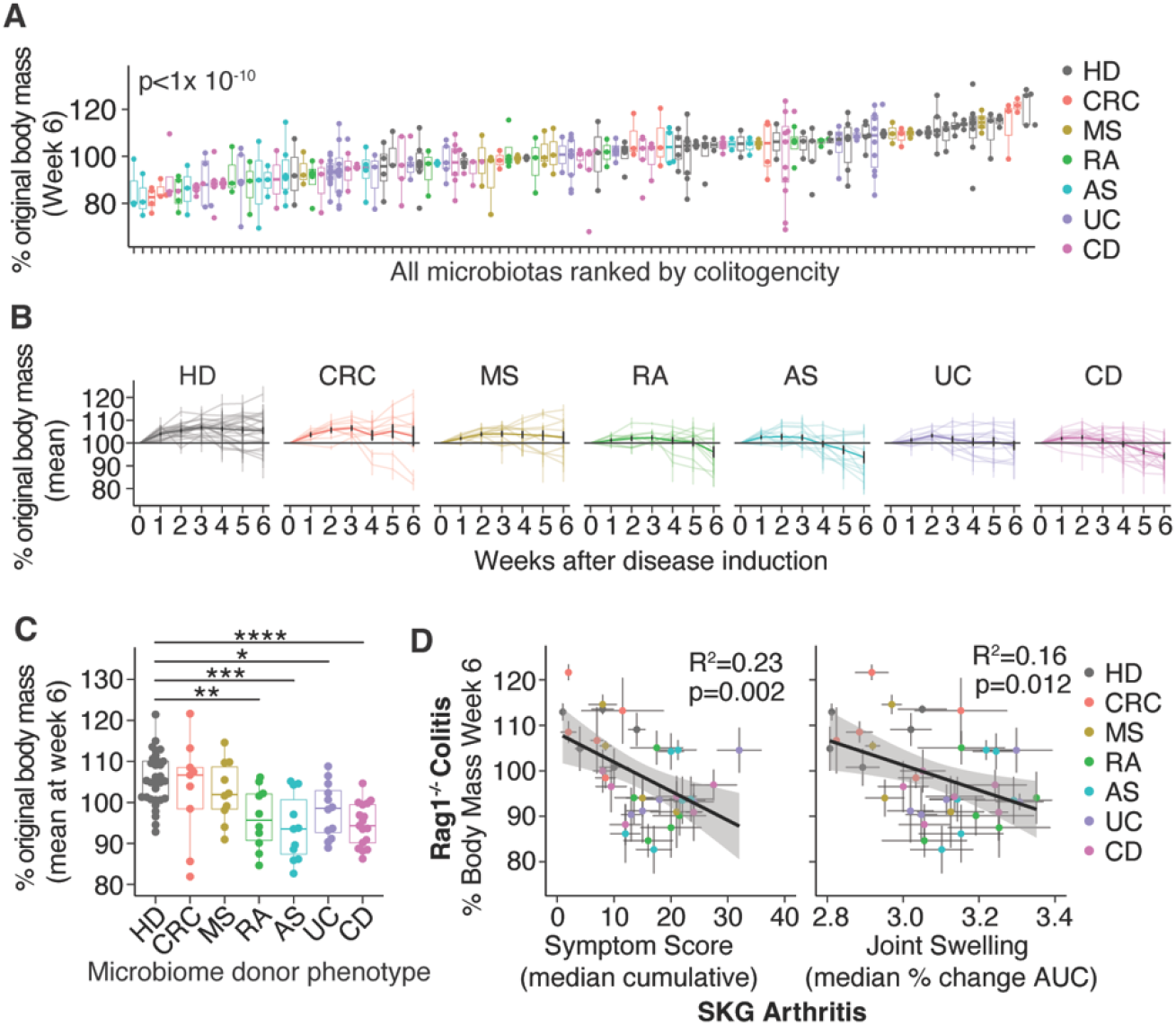
Microbiomes from humans with diverse IMID enhance colitis severity in susceptible mice. **(A)**, Median colitis severity six weeks after naïve T cell transfer (as assessed by loss in body mass) in groups of Rag1-/- mice colonized with one of 99 different human microbiomes. P value calculated by ANOVA. **(B)**, Change in body mass over time of groups of mice colonized with different human microbiomes. Lighter colored lines show mean +/- SEM change in body mass of mice colonized with each microbiome; darker lines show the weighted mean +/- SEM change in body mass of mice colonized with microbiomes from humans with the indicated disease. Grey horizontal lines indicate baseline body mass. **(C)**, The median change in body mass six weeks after naïve T cells transfer in groups of mice colonized with human microbiomes from donors with different heath statuses. Each point represents mean data of a group of mice colonized with a single microbiome. * p<0.05, ** p<0.01, ***p<0.001, ****p<0.0001 Dunnett’s test comparing disease groups to HD. **(D)**, A comparison of disease severity in the colitis model (change in body mass) and the SKG arthritis model (left panel; symptom score, right panel; joint swelling) in groups of mice colonized with the same human microbiomes. In panels A-C, N human microbiomes of each disease tested in n mice: AS, N=11, n=36; CD, N=17, n=75; CRC, N=9, n=29; HD, N=30, n=116; MS, N=10, n=29; RA, N=10, n=34; UC, N=11, n=72. In panel D, total microbiomes = 36. Boxplots show median +/- IQR in the style of Tukey.

### Inter-microbiome variation in colitis and arthritis severity is correlated

We compared the degree of disease severity in the Rag1^-/-^ colitis and SKG arthritis models when mice are colonized with the same human microbiomes. We observe a significant correlation between the degree of weight loss in the colitis model with both symptom scores and the degree of joint swelling in the SKG model (R^2^=0.23 and 0.16; p=0.002 and 0.012, respectively; Figure 3D). Change in body mass was also correlated between mice in each disease model when colonized with the same human microbiome (R^2^=0.21, p=0.003; Figure S4A). The correlation between disease in the colitis and arthritis model was not solely driven by the effect of exacerbated disease in mice colonized with AS, RA, CD and UC microbiotas. Comparing disease severity in the models only in mice colonized with HD, MS or CRC microbiomes (which experience relatively mild disease), there remains a correlation between weight loss in the colitis model and both symptom scores and weight loss in the SKG model (R^2^ = 0.4 and 0.2; p=0.009 and 0.06, respectively. Figure S4B). When colonized with the same microbiomes, disease severity in the EAE model was not correlated with colitis or arthritis severity (Figure S4C). HLA B27 is strongly associated with AS and has also been associated with changes in microbiome composition independent of an AS diagnosis (*44*). Within the donors included in the colitis model experiments, 9/10 AS donors were B27^+^. Although HLA status was unknown for the majority of healthy donors, we did include two healthy donors known to be B27^+^. It is notable that colitis severity was particularly mild in mice colonized with these B27^+^ healthy donors (Figure S4D), suggesting that B27 status alone does not explain enhanced colitogenicity of AS donor microbiomes.

### Colitis is more severe in mice colonized with microbiomes from humans with elevated CRP

To further explore patterns of function within microbiomes from human donors with varied diagnoses we clustered the microbiomes based on the severity of disease in the EAE, colitis and arthritis models. This segregated microbiomes by the diagnosis of the donor (Figure S5A, p=0.013, PERMANOVA). Notably, microbiomes from donors with MS clustered apart from microbiomes from healthy donors and donors with other IMID, whereas AS, RA, UC and CD microbiomes clustered together (Figure 4A and S5B; p=0.001, PERMANOVA). We considered that there are characteristics of IBD, AS and RA that set these diseases apart from MS. For example, the role TNFα-driven fibrotic tissue damage in IBD and inflammatory arthritis. Indeed, grouping these diseases based on FDA-approved immune therapeutic targets (as a proxy for immunological mechanism) produces a similar clustering to that which we see when clustering on the data from the microbiome transfer experiments (Figure 4B). Another way in which CD, UC, RA and AS manifest differently to MS is in systemic inflammatory markers. For example, CRP, erythrocyte sedimentation rate and neutrophil/lymphocyte ratio are frequently elevated relative to healthy controls in IBD, RA and AS but rarely so in MS (*45-48*).

**Fig. 4:**
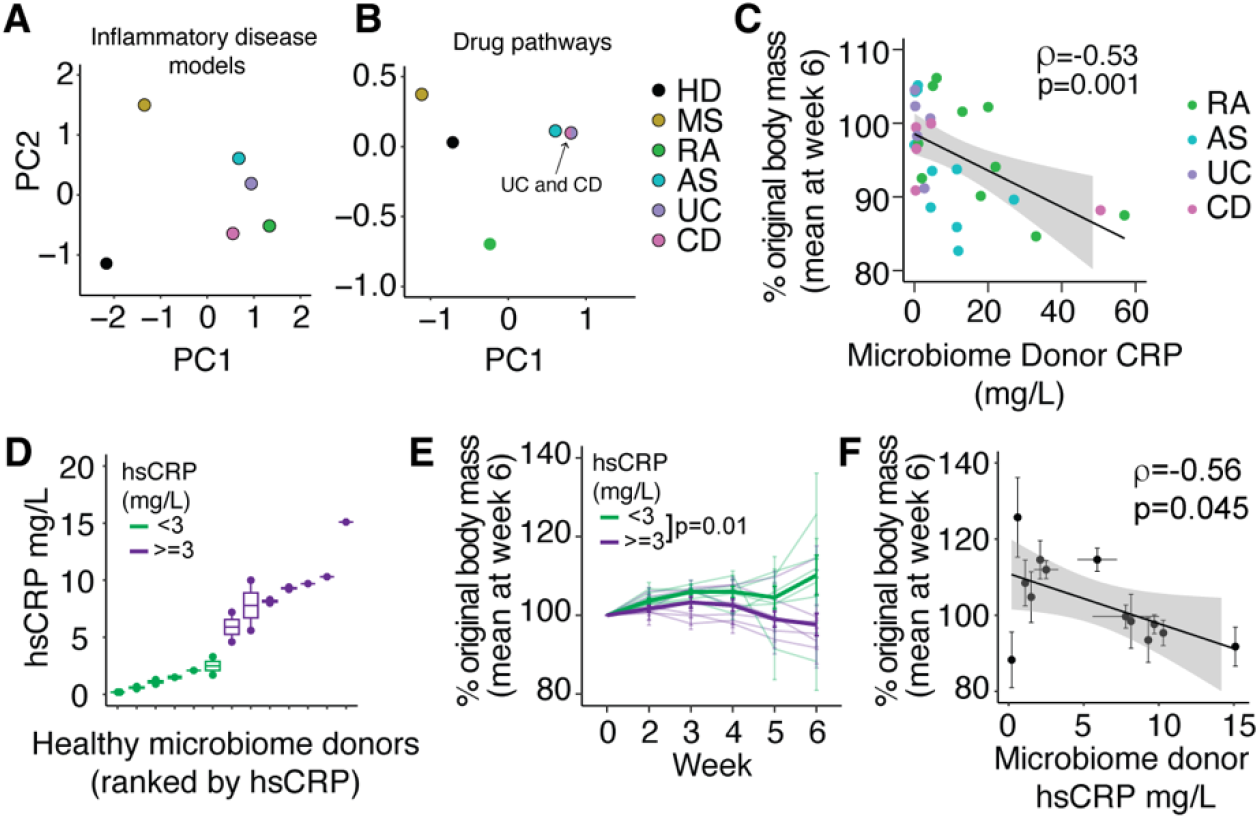
Colitis is more severe in mice colonized with microbiomes from humans with elevated serum CRP. (A)Principal component analysis of median EAE, arthritis and colitis severities of all microbiomes from each disease group (Euclidian distances). **(B)** Principal component analysis of FDA-approved immune therapeutic targets for each disease group (Jaccard). **(C)**, Correlation between microbiome donor serum CRP and colitis severity and groups of mice colonized with microbiome from those donors. **(D)**, Serum hsCRP values in a cohort of healthy subjects, including some with low-grade inflammation. **(E)**, Colitis severity (as measured by loss in body mass) in mice colonized with microbiomes from healthy humans with or without low-grade inflammation. **(F)**, The relationship between microbiome donor serum hsCRP and colitis severity in groups of mice colonized with microbiomes from those donors. Mean +/- SEM from groups of mice and mean hsCRP (where multiple values are available) are shown. r values show Pearson correlation coefficients. Panels E and F include data from 13 microbiomes assayed in a total of 78 mice.

C-reactive protein (CRP) is an acute-phase protein secreted by the liver in response to IL-6. It is a sensitive and dynamic biomarker of inflammation and is commonly used clinically to assess inflammation in the context of infection, injury and certain IMIDs (*49*). We found that the serum concentration of CRP of a human microbiome donor at the time of fecal microbiome collection is positively correlated with the severity of colitis in mice colonized with that microbiome (R^2^=0.32, p=0.005; Figure 4C). The association between donor serum CRP and colitis severity is also significant considering the RA and AS groups individually (R^2^=0.32 and 0.33 and p=0.05 and 0.05, respectively; Figure S5C). Colitis severity in mice colonized with human microbiomes was not significantly associated with other measures of human disease activity. This included Disease Activity Score (DAS28, not including CRP) for RA (*50*), Bath Ankylosing Spondylitis Disease Activity Index (BASDAI) for AS, endoscopic disease severity measures for IBD and fecal calprotectin concentrations for a subset of IBD and AS microbiome donors (Figure S5D). Focusing on mice colonized with microbiomes from donors with MS, we found no correlation between colitis or arthritis severity and patient disease severity (Extended Disability Severity Score) or disease duration (Fig S5E).

Although most commonly associated with overt pathology, a minority of healthy individuals exhibit persistently elevated CRP in the absence of any specific infection, injury or IMID diagnosis. Defined as a concentration of CRP between 3 and 10 mg/L (as measured by a high sensitivity [hsCRP] assay), this phenomenon is often referred to as low-grade inflammation and is a significant risk factor for a range of morbidities, most notably cardiovascular and metabolic disease (*52*). To further investigate the link between serum CRP and microbiome-driven inflammatory responses we assembled a cohort of 13 individuals who exhibited a range of hsCRP levels despite being clinically healthy; specifically having no diagnosis of infection, cancer, CD, UC, inflammatory arthritis, MS, type-1 or type-2 diabetes, cardiovascular disease or systemic lupus erythematosus (Figure 4D). These donors are not included in prior analyses. For eight of the 13 donors, we obtained two samples at least four weeks apart, and we observed stable levels of hsCRP in these individuals (Figure 4D and S6A). LGI is often observed during aging and in individuals who are obese so we took care to match donor age and BMI in the low hsCRP and elevated hsCRP groups (Figure S6B). We colonized groups of germ-free Rag1^-/-^ mice with the fecal microbiomes from these 13 donors and assessed colitis severity following naïve CD4 T cell transfer. Microbiomes from humans with LGI transferred more severe colitis to the mice compared to microbiomes from humans with hsCRP under 3 mg/L (Figure 4E). Furthermore, microbiome donor hsCRP concentration was correlated with colitis severity (p=0.045; Figure 4F). Importantly, colitis severity was not correlated with the common LGI risk factors age or BMI (Figure S6C).

## DISCUSSION

Using three mouse models of inflammatory disease, we report a comprehensive assessment of the inflammatory potential of the human microbiome, including microbiomes from individuals across a spectrum of immune-mediated inflammatory diseases and inflammation-associated cancer.

Our analysis of the microbiomes in MS patients and their role in the EAE mouse model reveals two important findings. First, the microbiomes from MS patients have unique effects that differ from those of healthy donors and from microbiomes associated with other IMIDs. This suggests that the microbiome of MS patients has specific changes that are not seen in other IMIDs. Second, when we examine each microbiome individually and as a group specific to MS, their effects in the EAE model are distinct compared to their effects in models of colitis or arthritis. This demonstrates that the influence of microbiome on EAE severity is through a different mechanism than in the colitis and arthritis models. A clear distinction between EAE and the other models is the requirement for leukocytes to cross the blood brain barrier, the integrity of which can be altered by the gut microbiome (*53, 54*). The microbiome can also influence the function of astrocytes and microglia, and modulation of these CNS-specific cellular processes could underlie the EAE-restricted effects we observe (*55, 56*).

We included microbiome samples from a cohort of individuals with treatment-naïve CRC. Although cancer cannot strictly be considered immune-mediated, several forms of cancer are strongly linked to inflammation. This is typified by the increased risk of CRC in individuals with ulcerative colitis. In the three tested mouse models, microbiomes from donors with CRC behave on average as microbiomes from healthy donors. We conclude from this that the inflammatory potential of these microbiomes is not intrinsically altered. This does not preclude a role for the microbiome in the pathogenesis of CRC, as it is known that the microbiome can contribute to carcinogenesis through provision of genotoxic agents and pro-proliferative signals (*57*). Furthermore, we did not include microbiomes from donors with colitis-associated CRC, a subgroup where the role of microbiome-driven inflammation may be especially relevant.

We find a strong relationship between the effect of each human microbiome on colitis in Rag1^-/-^ mice and arthritis in SKG mice. We also find that the influence of microbiomes from humans with IBD and inflammatory arthropathies are similar in all mouse models. Clustering microbiomes on the basis of disease severity in the three models segregates AS, RA, CD and UC from healthy donors and highlights the distinct response we observe in mice colonized with microbiomes from donors with MS. There are several parallels between the underlying pathology and therapy of these diseases, both in human patients and in mouse models. For example, the immune landscape of lesional tissue is dominated by Th1 and Th17-polarized T cells with inflammatory fibroblasts, monocytes and macrophages (*58, 59*). Anti-TNFα and Jak/Stat inhibitors are mainstays of therapy for both gastrointestinal and rheumatological IMIDs. Multiple sclerosis is also distinguished from these other IMID by a lack of systemic inflammatory markers. For example, elevated CRP, erythrocyte sedimentation rate and neutrophil/lymphocyte ratio are common in IBD, RA and AS but rare in multiple sclerosis (*45-48*).

We further examined the relationship between systemic inflammation and microbiome function, finding a correlation between microbiome donor serum CRP and the inflammatory potential of the microbiome.

Correlations between serum CRP and microbiome composition have been identified in numerous IMIDs, including in AS (*60, 61*) and CD (*62, 63*). In a large metagenomic study of microbiomes from patients with AS and RA, serum CRP levels explained a significant amount of variation in microbiome composition (*5*). In a cohort of individuals with mood disorders (but no overt IMID), elevated hsCRP was associated with changes in microbiome composition and the fecal metabolome (*64*). These data strongly suggest a link between systemic inflammation in the host and microbiome composition. The directionality of this interaction is unknown, but we hypothesize that interactions between microbiome and the host immune system, particularly in the context of a compromised epithelial barrier (i.e. ‘leaky gut’), contributes to the development of LGI. A study in an aging population found a correlation between serum hsCRP and LPS-binding protein, a marker of intestinal barrier dysfunction (*65*). This supports a role for epithelial barrier defects and the microbiome in LGI. Furthermore, induction of dysbiosis with microbiome-depleting antibiotics modulates hsCRP levels, including in healthy human volunteers (*66*). It is also possible that host inflammation arising independent of the microbiome could lead to changes in microbiome composition. For example, the microbiome of IL-10-deficient mice is altered as colitis develops, becoming more colitogenic when transferred to new recipients (*67*).

In summary, we identify cross-disease patterns of microbiome pathogenic functionality that parallel other shared characteristics between IBD and inflammatory arthritis including pathological mechanisms and therapeutic targets. We identify a link between systemic inflammation and pathogenic microbiome function that exists across IMID and in humans with low-grade inflammation.

## MATERIALS AND METHODS

### Human subjects

#### Multiple Sclerosis

Human fecal samples were collected from ten donors with MS recruited from the Corinne Goldsmith Dickinson Center for MS at Mount Sinai Hospital (New York, NY). The study was approved by Mount Sinai’s Institutional Review Board and all participants provided written informed consent. Six were male, four were female, all self-identified as White non-Hispanic and had a median age of 49 (+/1 16.9; SD). Of the ten donors, seven were diagnosed with relapsing remitting MS, two with primary progressive MS and one with clinically isolated syndrome.

Inclusion criteria required that participants carry a diagnosis of MS, be of White (Hispanic or non-Hispanic) ethnicity. Exclusion criteria included the presence of other autoimmune disorders, gastrointestinal infections, and other neurological disorders. Participants were excluded if they received oral antibiotics within the past three months, corticosteroids within the past 30 days, or were on a disease-modifying therapy for less than three months.

#### Rheumatoid arthritis and ankylosing spondylitis

Fecal samples from donors with AS and RA and associated controls were collected as part of the International Arthritis Microbiome Consortium (IAMC) (*5*), which collated samples from patients in Oxford, UK (Nuffield Orthopedic Center and Oxford biobank; the primary source of AS samples), Birmingham, UK (the Birmingham Early Arthritis Cohort from Sandwell and West Birmingham NHS Trust and University Hospitals Birmingham NHS Foundation Trust; the primary sources of RA samples). This study was reviewed and approved by local Research Ethics Committees, specifically, in Oxfordshire (REC 06/Q1606/139), and the West Midlands-Black Country (REC 12/WM/0258). The median age of AS subjects was 50 +/-14 years. One subject was female, ten were male. Nine of the eleven AS donors were HLA B27 positive, one negative and one unknown. Individuals with AS frequently experience complex comorbidities and this was reflected in this cohort. Three AS donors were additionally diagnosed with uveitis, three with peripheral arthritis, one with dactylitis, one with psoriasis and one with IBD (unspecified). Of the donors with RA, five were male and five female, the median age was 65.5 +/-13.1 years. All RA subjects were rheumatoid factor and anti-cyclic citrullinated peptide (CCP) positive.

#### Inflammatory bowel disease

Fecal samples from donors with IBD (UC or CD) and control samples were obtained from participants in one of three previously described US-based studies; the Mount Sinai Crohn’s and Colitis Registry (MSCCR) cross-sectional study (*68, 69*) (Mount Sinai Hospital IRB: 11-01669, the Road-to-Prevention study (Mount Sinai Hospital IRB: 16-00021) (*70*), which both collected samples from donors in the New York area, and a cohort established at Cedars Sinai Medical Center (Cedars-Sinai Medical Center Institutional Review Board IRB 3766) (*71*) which collected samples from the Los Angeles area.

#### Colorectal cancer

Fecal samples from donors with early-stage primary tumors in the colon or rectum were obtained from a contract research organization (BioIVT). Samples were collected before chemotherapy or surgery. No donor had taken antibiotics in the three months before sample collection. Of the nine donors used in this study, seven were diagnosed with cancer of the colon and two with rectal cancer. All donors were assessed as having tumor grade 2 disease, six with T3N0M0 disease and three with T3N0M1 disease. The average age of these donors was 65 +/-6 years. Five were male, four female and all self-identified as Caucasian. These donors had no diagnosis of autoimmune or inflammatory disease, renal or liver disease, dementia, significant mental illness or non-CRC hematological/oncological disease.

#### Healthy controls

Healthy control samples (from donors with no diagnosis of chronic infection, inflammatory or immune-mediated disease or cancer) were obtained from controls in five different IMID cohorts in the USA and the UK. Of a total of thirty healthy donors, three were collected alongside samples from the MS cohort, six controls were from the Mount Sinai Crohn’s and colitis cohort, five controls were from the IAMC (RA and AS) cohort and nine controls from the RTP Crohn’s and colitis cohort.

#### Low-grade inflammation

Fecal samples from healthy donors with paired contemporaneous serum hsCRP measurements were obtained from a contract research organization (BioIVT) and from the MSCCR cohort (*68, 69*) (Mount Sinai Hospital IRB: 11-01669). These donors had no diagnosis of infection, cancer, CD, UC, inflammatory arthritis, MS, type-1 or type-2 diabetes, cardiovascular disease or systemic lupus erythematosus. No donor had taken antibiotics in the three months before sample collection. Of these donors, four were female, nine male and their median age was 53+/-18 years. Two donors self-identified as Black or African-America, three as Hispanic, seven as White or Caucasian (one unknown race/ethnicity).

### Human fecal microbiome sampling

Human fecal samples were frozen and stored at -20^°^C before homogenization under strict anaerobic conditions in pre-reduced LYBHI media (37 g/L BD Brain Heart Infusion, 5 g/L yeast extract, 1 g/L D-xylose, D-fructose, D-galactose, cellubiose, maltose and sucrose, 0.5 g/L N-acetylglucosamine and L-arabinose, 0.5 g/L L-cysteine, 1 g/L malic acid, 2 g/L sodium sulfate, 0.05% Tween 80, 20 μg/mL menadione, 5 mg/L hemin, 0.1 M MOPS, pH 7.2) at approximately 50 mg/mL. Samples were cryopreserved at -80^°^C at approximately 2.5 mg/mL in LYBHI media supplemented with 15% glycerol. The CFU of 51 of the samples (including samples from all cohorts and all disease states) was estimated by plating on non-selective rich media (BD BBL Chocolate II Agar) under anaerobic conditions. The mean CFU/g of these samples was 7.1 × 10^8^ (range 4.3 × 10^6^ – 4.8 × 10^9^) and there was no significant difference in the CFU of samples from donors with different diseases (p=0.1, ANOVA) or from different study cohorts (p=0.47, ANOVA).

### Mouse breeding and husbandry

Germ free C57Bl/6J, Rag1^-/-^ on the C57Bl/6J background and SKG mice (*39*) (harboring the W163C ZAP-70 mutation on the BALB/c background, a gift of R. Thomas) were bred in flexible vinyl isolators at the Icahn School of Medicine gnotobiotic facility. All mice were provided autoclaved chow (5K67, Labdiet) and water *ad libitum*. The germ-free status of each colony was regularly confirmed by a combination of molecular and microbiological methods. At 4-5 weeks of age, germ-free mice were colonized with human fecal material by a single gavage with approximately 0.5 mg of homogenized fecal material in 200 μl reduced LYBHI media. This equated to administering approximately 3.6 × 10^6^ +/-5.8 × 10^6^ CFU per mouse (mean +/- SD). From the time of colonization, mice were housed in individually-ventilated, positive pressure cages (Allentown) with autoclaved 5K67 (Labdiet) chow and water and handled using strict aseptic methods. For all experiments, disease was induced in mice 2-3 weeks after colonization. An approximately equal number of male and female mice were used.

### Naïve T cell transfer-induced colitis in Rag1^-/-^ mice

CD4^+^ T cells were isolated from the spleens of specific-pathogen-free C57Bl/6J mice (Jackson Labs) using negative magnetic enrichment (MagniSort Mouse CD4 T cell Enrichment Kit, Thermofisher Scientific). The resulting CD4^+^ T cells were immunolabelled with anti-mouse CD4 (GK1.5, PerCP-Cy5.5), CD25 (PC61, PE) and CD45RB (C363-16A, FITC) and naïve T cells were enriched using a FACS BD ARIA II as CD4^+^, CD25^-^ and CD45RB^HI^. Purity of the desired population routinely exceeded 97%. Three weeks after colonization, Rag1^-/-^ mice received 1 × 10^6^ sex-matched naive CD4 T cells by intraperitoneal injection. To enable the scale of experiments required for these studies, body mass was used as the primary measure of disease severity for all mice. Although we have previously found that body mass correlates with immune and histologic markers of colitis severity in this model, these mice can develop systemic inflammation (resulting in loss in body mass) that can be functionally uncoupled from colitis as they are mediated by different cytokine pathways (*72*). Therefore, for a substantial subset of mice we assessed levels of fecal LCN2 and colon histopathology as direct measures of intestinal inflammation. Both of these measures were significantly correlated with loss in body mass (Figure S3E). Any animal presenting with weight loss greater than 80% was euthanized and for analysis the final recorded weight was carried forward to any subsequent remaining timepoints.

### Arthritis induction in SKG mice

Disease was induced in SKG mice three weeks after colonization by intraperitoneal injection of 3 mg of curdlan ((1,3)-β-glucan; Waco) in 300 μl of sterile PBS. Disease was monitored by measuring weight loss, joint swelling measured using calipers (the dorsal-palmer dimension of front paw, dorsal-plantar dimension of hind paw footpad, and width of hind paw) and a symptom scoring system: 0; no swelling or redness, 0.5; very mild swelling or redness of digits and redness of the joint, 1; mild swelling of the joint, 2; moderate swelling the joint, 3 substantial swelling of the joint. The sum of the score for all four paws is reported. In the absence of joint symptoms but the presence of other early disease manifestations (e.g. reddening or swelling of the skin at the ears and snout which commonly preceded joint symptoms) a total score of 2 is given. Any animal presenting with a score of >11 for 2 consecutive days or weight loss >20% was euthanized and for analysis the final recorded score, joint dimensions and weight were carried forward to any subsequent timepoints.

### EAE induction in C57Bl/6 mice

Complete Freund’s adjuvant (BD Difco), supplemented with 4 mg/mL killed Mycobacterium tuberculosis H37Ra (BD Difco) was emulsified with an equal volume of 3 mg/mL myelin oligodendrocyte glycoprotein (MOG) residues 35-55 (GL Biochem) in sterile PBS. Three weeks after colonization, mice were injected with 100 ml of this emulsion (150 mg MOG 35-55 per mouse) subcutaneously at the base of the tail. Mice also received 200 ng Pertussis toxin (Sigma) in 500 ml sterile PBS by intraperitoneal injection at the time of CFA injection and 48 hours later. Ascending paralytic disease was monitored using a scoring system: 0; no symptoms, 0.5; signs of ataxia or tail weakness, 1; tail paralysis, 1.5; hind limb ataxia or one-sided mild weakness 2; bilateral hind limb weakness and/or impaired righting reflex, 2.5; single-sided hind limb paralysis and absent righting reflex, 3; bilateral hind limb paralysis, 3.5; fore limb weakness, 4; forelimb paralysis, 5; dead or moribund. Any animal presenting with a score of 4 for two consecutive days or with weight loss >20% was euthanized and afforded a score of 5 for that day and any subsequent days.

### Histology

Mouse colons were gently flushed with cold PBS, opened longitudinally and rolled before fixation in 10% buffered formalin for a minimum of 24 hours. Fixed tissue was stored in 70% ethanol. 4 micron sections of paraffin embedded tissue, cut such that the entire colon length is visible in one section, were stained with hematoxylin and eosin. Embedding, sectioning, staining and imaging were performed by HistoWiz according to standard protocols. Digitized images were scored based on the protocol described in Buonocore et al. (*73*).

Briefly, areas of the tissue representing the proximal, medial and distal regions of each tissue were scored using a semiquantitative scale from 0-4 based on the following criteria. 0: normal tissue appearance, 1: some evidence of epithelial hyperplasia without obvious immune infiltrates, 2: pronounced epithelial hyperplasia and obvious immune infiltrates with largely normal goblet cell appearance, 3: severe hyperplasia and immune infiltration with obvious loss of goblet cells, 4: as ‘3’ with the appearance of gross tissue damage in the form of transmural inflammation, ulceration and/or crypt abscesses. The scores from each three tissue regions were summed to give a total tissue score 0-12.

### Mouse fecal Lipocalin-2 measurements

Mouse fecal pellets were frozen at -20^°^C until processing. Fecal samples were weighed and homogenized at between 100 and 200 mg/mL in sterile PBS supplemented with 0.1% Tween-20 in a beadbeater (without addition of beads). Homogenates were then cleared by centrifugation and supernatants stored at -20^°^C until assayed.

Thawed fecal supernatants were diluted to 0.1 mg/mL (according to original fecal sample mass) in PBS supplemented with 1% BSA before lipocalin-2 concentrations were assayed by ELISA (R&D Mouse Lipocalin-2/NGAL DuoSet) according to the manufacturer’s instructions.

### Human subject laboratory testing

Calprotectin was measured in fecal samples by ELISA (BUHLMANN EK-Cal) according to manufacturer’s instructions. Serum CRP concentrations of IMID subjects were measured as part of standard clinical testing. hsCRP measurements in healthy subjects reported in Fig 4 and S4 were performed by Quest Diagnostics.

### Statistical analysis

All statistical analysis and plotting were performed using R v4.1.2. Plots were constructed using ggplot2 v3.5.1 and ggbeeswarm v0.7.2. When comparing multiple groups to a control group, DescTools v0.99.55 (*74*) was used to perform Dunnett’s tests. The cor.test and aov functions in stats v3.6.2 were used to calculate Pearson correlation coefficients and perform ANOVA analyses, respectively. Wilcoxon tests were performed using the stat_compare_means function of ggpubr v0.6.0. In Fig 4F, a pair of time courses are compared and the lme function of nlme v3.1was used to produce a linear mixed effects model. PCA and PERMANOVA analysis was performed using the vegdist, prcomp and adonis2 functions in vegan v2.6. A minority of the colitis model data was previously included in analyses published in Britton et al. 2019 (*38*). This includes data from 9 of the 30 HD, 5 of the 12 UC and 6 of the 17 CD microbiomes included in this manuscript. A breakdown of the previously published and newly generated colitis data for HD, UC and CD microbiomes is shown in Figure S7. In some instances, the data presented in this manuscript is a pool of data included in the previous publication and additional experimental replicates using the same microbiome.

## Acknowledgements

We thank G.N. Escano, C. Fermin, E. Vasquez, J. Eggers, and Z. Li for technical assistance and Ms A. Batki for support with patient recruitment. This work was performed with the support of staff and facilities of the Icahn School of Medicine Gnotobiotic Facility, the Microbiome Translation Center at Mount Sinai and the Mount Sinai Flow Cytometry Core. We thank Dr R. Thomas, University of Queensland, for the SKG mice and all investigators of the IAMC, the Road-to-Prevention study group and the Mount Sinai Crohn’s and Colitis Registry. We acknowledge the following sources of funding: Crohn’s and Colitis Foundation award 988073 (GJB), The Scherr Family Foundation and the Friedman Brain Institute at Mount Sinai (GJB, SKT), Versus Arthritis grant 21226 (funding for the IAMC), Crohn’s and Colitis Foundation awards 632758 and 651867 (JJF), National Institutes of Health NIDDK DK112978, NIDDK DK124133, NIDDK DK123749 (JJF), National Institutes of Health NIDDK DK131862 F30 (AC), Janssen Human Microbiome Institute (JJF), U.S. Department of Veterans Affairs Career Development Award IK2CX001717 (JPJ), UK National Institute for Health Research Oxford Biomedical Research Centre (PB), UK National Institute for Health Research Birmingham Biomedical Research Centre (KR), The Department of Neurology at Mount Sinai (SKT), UK National Institute for Health Research Oxford Biomedical Research Centre (PB). The views expressed are those of the authors and not necessarily those of the NIHR, the UK Department of Health and Social Care or any other funder.

## Author contributions

Conceptualization: GJB, JJF, Investigation: GJB, LB, IM, AC, TP, DH, GB, PC, Formal analysis: GJB, LB, Visualization: GJB, LB Methodology: GJB, IM, GB, IB, LHL, SJB, FK, PRM, Resources: LHL, SJB, JB, JJ, AND, SS, AF, PCT, PB, CH, DRL, MCD, KR, SKT, Funding acquisition, GJB, AGP, PCT, PB, CH, DRL, MCD, KR, SKT, JJF, Project administration: GJB, IM, PC, JJF, Supervision: JJF, Writing – original draft: GJB, Writing – review & editing: GJB, CH, KR, SKT, JJF. All authors approved the manuscript before submission.

## Data and materials availability

All data used to construct figures are available in the supplementary materials. Materials used in this study are available upon request to the corresponding author. Material transfer agreements may be required.

---

**Fig S1:**
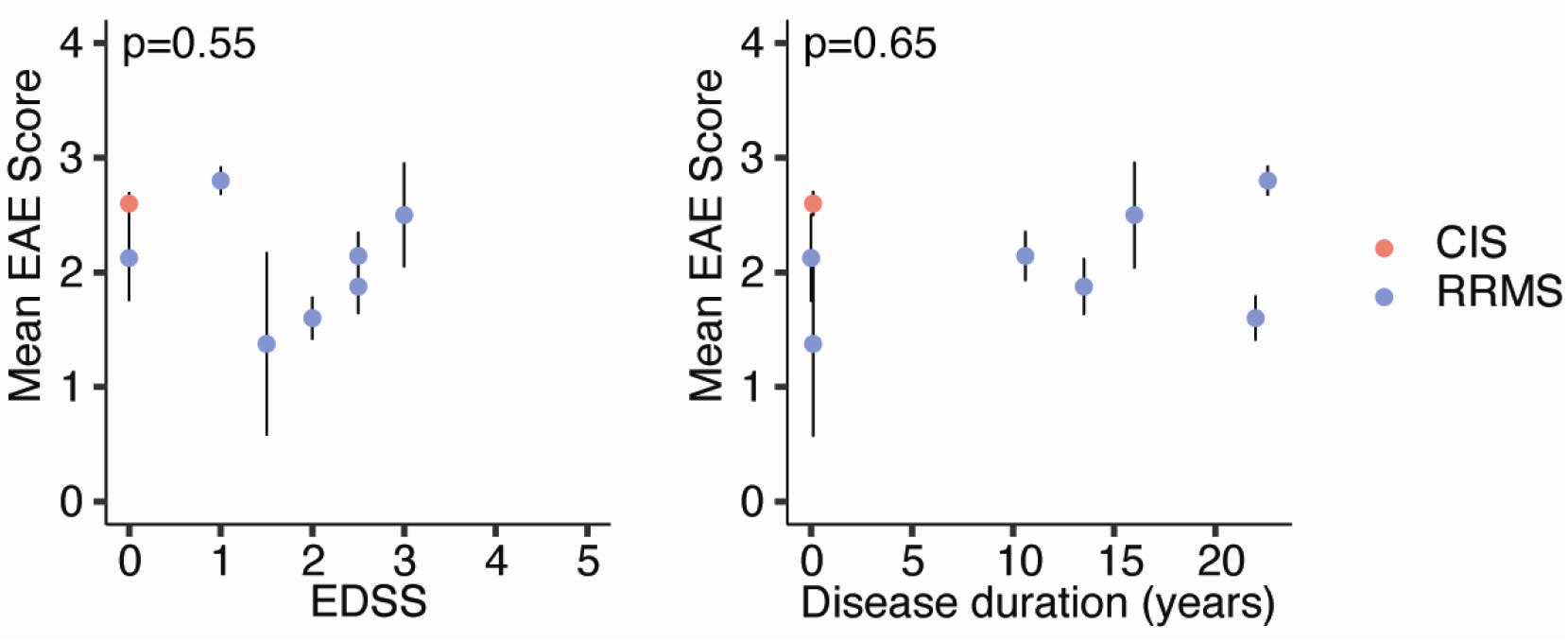
EAE severity in mice colonized with human microbiomes is not correlated with **(A)**, the Extended Disability Status Score (EDSS) or **(B)**, the time since diagnosis (disease duration) of the microbiome donor. RRMS – relapsing remitting MS, CIS – clinically isolated syndrome.

**Fig S2:**
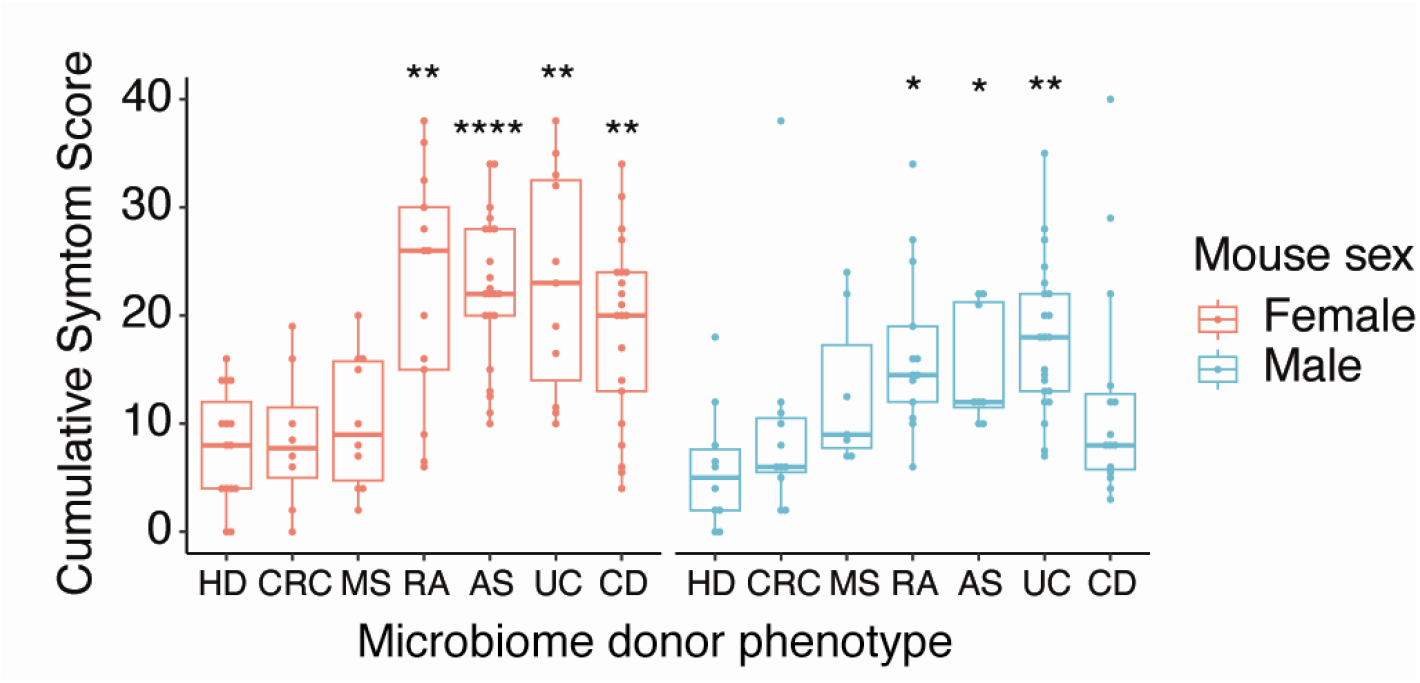
Arthritis severity (cumulative symptom scores) in SKG mice colonized with human microbiomes from donors of the indicated disease state. Each point shows data from one mouse. Box plots show median +/- IQR in the style of Tukey. * p<0.05, ** p<0.01, ***p<0.001, ****p<0.0001 Dunnett test comparing disease group to HD group, treating males and females as individual datasets. (legend over page)

**Fig S3:**
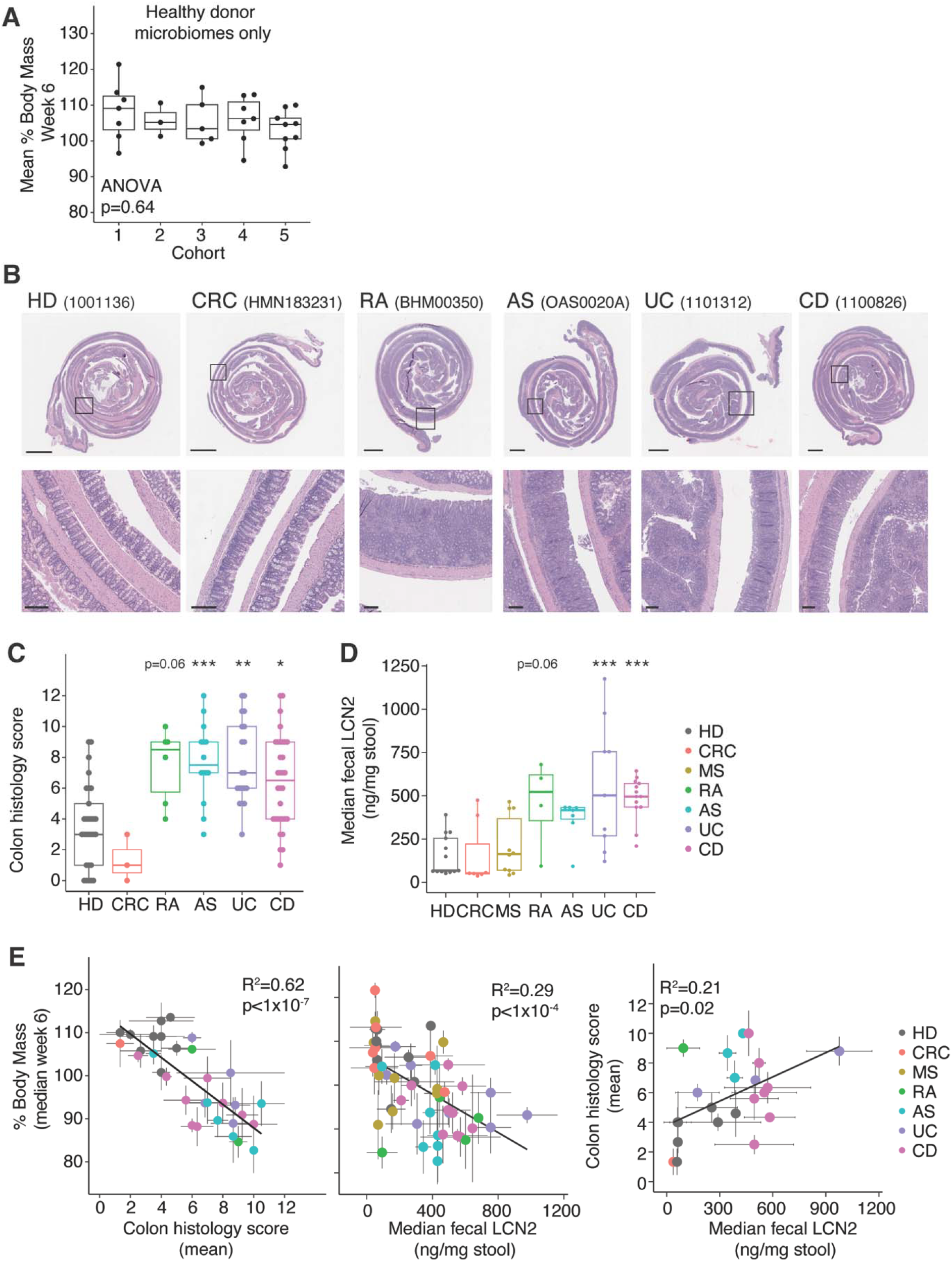
**(A)**, Colitis severity (median change in body mass six weeks after naïve T cells transfer) in groups of Rag1^-/-^ mice colonized with human microbiomes from healthy donors from five different cohorts. Each point represents mean data of a group of mice colonized with a single microbiome. **(B)** Representative photomicrographs of H&E-stained sections of colon tissue from mice colonized with the indicated microbiota 6 weeks after naïve T cell transfer. Scale bars in the upper panels represent 2mm, in the lower panels 200μm. **(C)** Semi-quantitative scoring of histopathology in mice colonized with microbiotas from humans of the indicated disease state 6 weeks after T cell transfer. Each point represents the score from a single animal. **(D)** LCN2 concentrations in fecal samples collected 6 weeks after T cell transfer in mice colonized with microbiotas from humans with the indicated disease. Each point represents the median LCN2 concentration from a group of mice colonized with a single human microbiome. **(E)** Correlations between changes in body mass, colon histopathology and fecal LCN2 concentrations in mice colonized with the same microbiomes 6 weeks after T cell transfer. Each point represents mean or median data f(as indicated) rom a group of mice colonized with a single human microbiome +/- SEM. Box plots show median +/- IQR in the style of Tukey. * p<0.05, ** p<0.01, ***p<0.001 Dunnett test comparing disease group to HD group.

**Fig S4:**
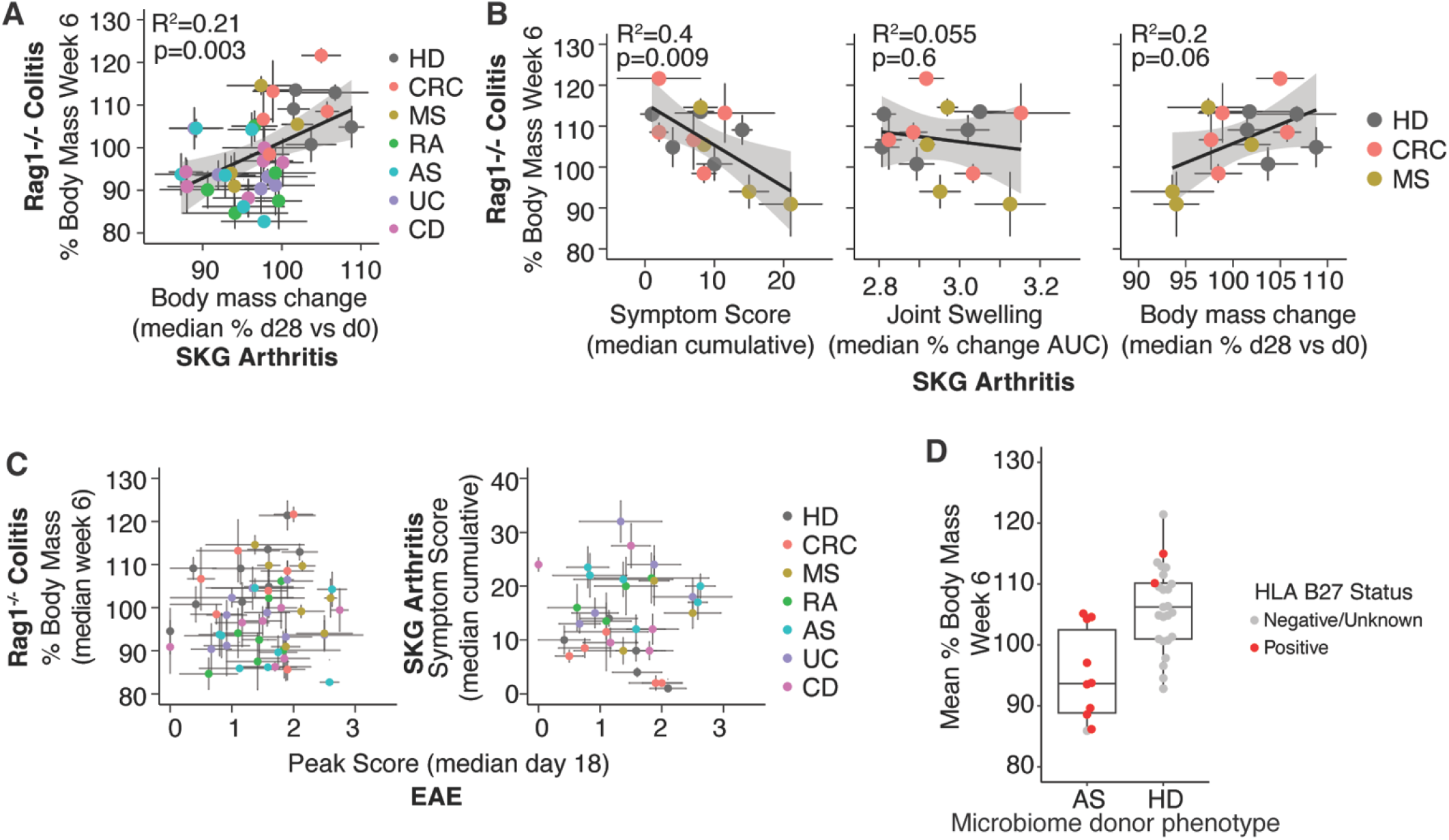
**(A)**, A comparison of change in body mass in the colitis model and the SKG arthritis model in groups of mice colonized with the same human microbiomes. Each point represents mean data of a group of mice colonized with a single microbiome +/- SEM. **(B)**, A comparison of disease severity in the colitis model (change in body mass) and the SKG arthritis model (left panel; symptom score, center panel; joint swelling, right panel, body mass) in groups of mice colonized with the same human microbiomes, including only MS, CRC and healthy donor microbiomes. Each point represents mean data of a group of mice colonized with a single microbiome +/- SEM. **(C)**, A comparison of disease severity in the EAE model (median peak disease score) and disease in the colitis model (change in body mass; left panel) and the SKG arthritis model (right panel; symptom score) in groups of mice colonized with the same human microbiomes. Each point represents mean data of a group of mice colonized with a single microbiome +/- SEM. **(D)**, Colitis severity (median change in body mass six weeks after naïve T cells transfer) in groups of Rag1^-/-^ mice colonized with human microbiomes from healthy donors or donors with AS, indicating the B27 status of the microbiome donor. Boxplots show median +/- IQR in the style of Tukey.

**Fig S5:**
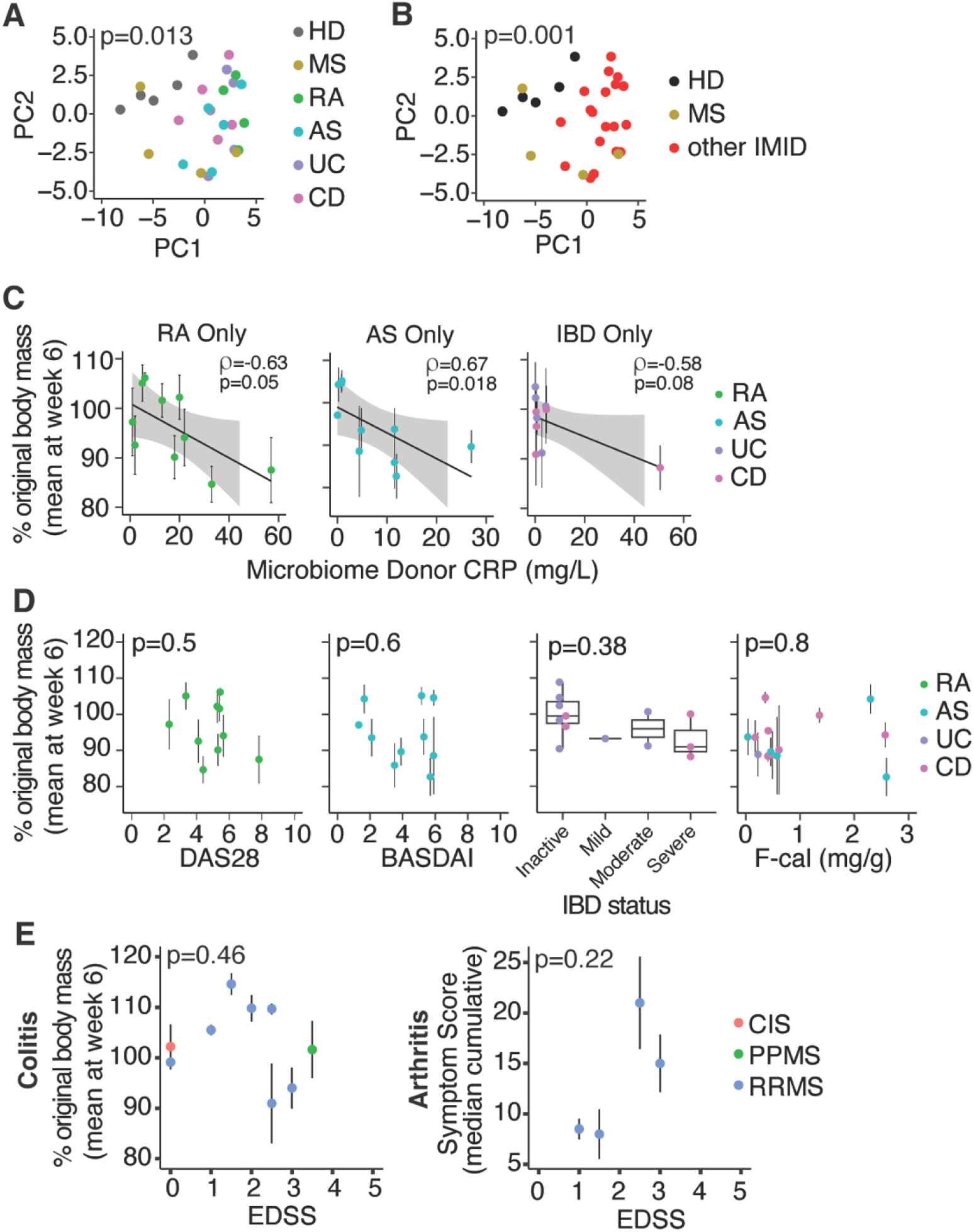
**(A, B)** Principal component analysis of a similarity matrix based on EAE (median cumulative disease score), arthritis (median cumulative disease score) and colitis (mean weight loss week 6) severities for microbiomes that were tested in all models colored by microbiome donor disease. P value calculated by PERMANOVA. **(C)** Correlation between microbiome donor serum CRP and colitis severity in groups of mice colonized with microbiome from those donors. **(D)**, Microbiome donor IMID severity (DAS28 exclusive of CRP for RA and BASDAI for AS, endoscopic measures for IBD and fecal calprotectin concentrations for a subset of IBD and As donors) compared to mean colitis severity in groups of mice colonized with each donor’s microbiome. IBD severity P value calculated by ANOVA. **(E)**, MS microbiome donor disease severity (Extended Disability Severity Score; EDSS) compared to mean colitis severity and median arthritis severity in groups of mice colonized with each donor’s microbiome.

**Fig S6.**
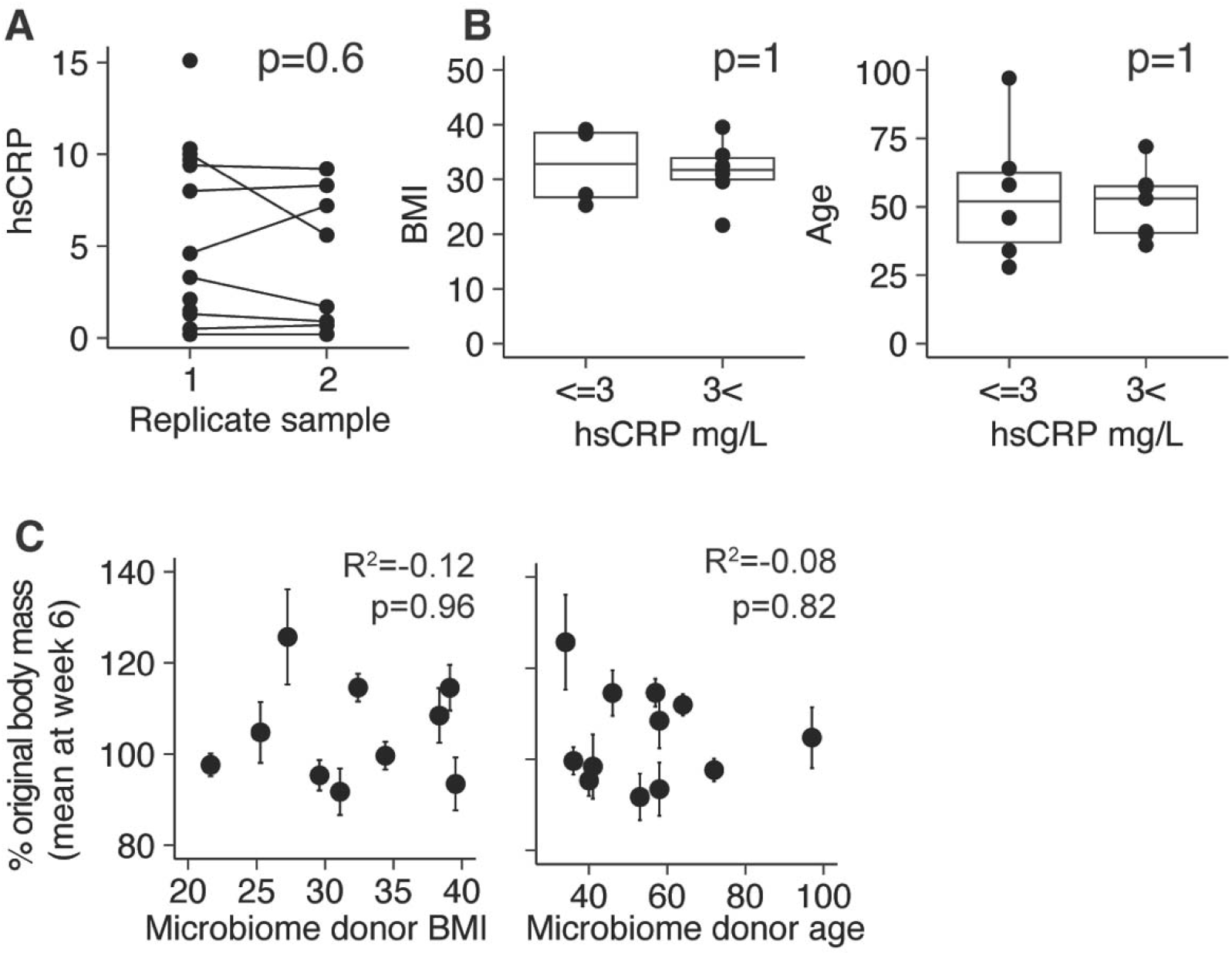
**(A)**, Serum hsCRP levels in microbiome donors, including 8 donors with replicate samples. Groups are compared with a paired t-test. **(B)**, Body mass index of microbiome donors stratified according to elevated (>3 mg/L) unelevated (=<3 mg/L) hsCRP and age (in years) of microbiome donors stratified according to elevated (>3 mg/L) unelevated (=<3 mg/L) hsCRP. **(C)**, Colitis severity (median change in body mass six weeks after naïve T cells transfer) in groups of Rag1^-/-^ mice colonized with human microbiomes is not correlated with donor BMI or age. Boxplots show median +/- IQR in the style of Tukey.

**Fig S7:**
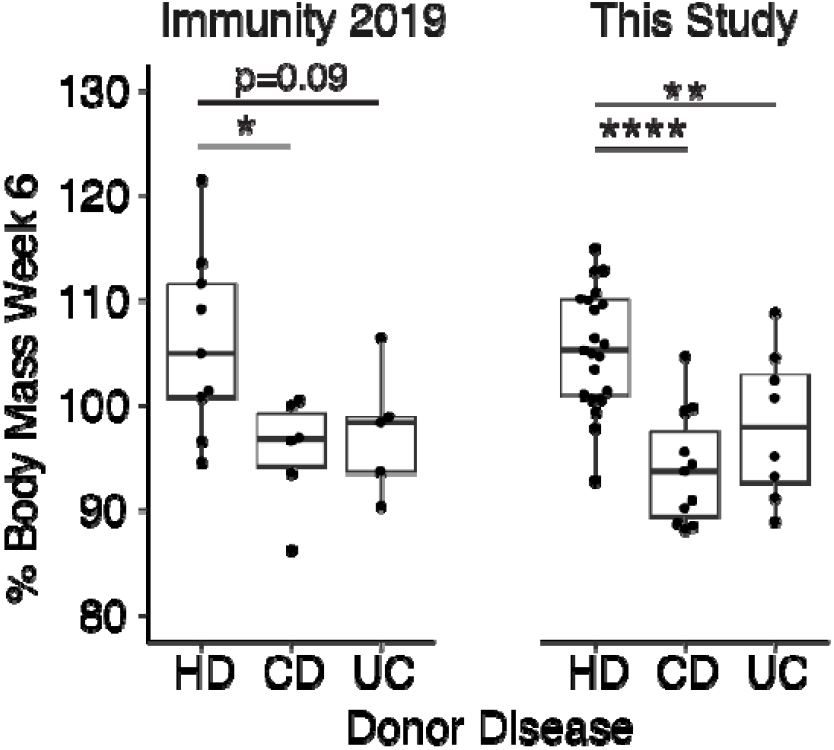
A breakdown of the subset of colitis data that was previously published in Britton et al. 2019 (*38*). The ‘Immunity 2019’ group includes all colitis data from mice colonized with fecal microbiomes that was analyzed in Britton et al. 2019 pooled with any additional replicates of those microbiomes that were subsequently performed. The ‘This study’ group shows only data from HD, UC or CD microbiomes that did not feature in our previous publication. For all analyses in this manuscript, the data from both groups are combined. Each point shows mean data from a group of mice. Boxplots show median +/- IQR in the style of Tukey. * p<0.05, ** p<0.01, ****p<0.0001, Dunnett test comparing disease group to HD group.

